# AFF3 and BACH2 are master regulators of metabolic inflexibility, β/α-cell transition, and dedifferentiation in type 2 diabetes

**DOI:** 10.1101/768135

**Authors:** Jinsook Son, Hongxu Ding, Domenico Accii, Andrea Califano

## Abstract

Type 2 Diabetes is associated with defective insulin secretion, reduced β-cell mass, and increased glucagon production. Cell lineage-tracing in rodents and human autopsy surveys support the notion of β-cell dedifferentiation as a unifying mechanism for these abnormalities. Yet, mechanistic determinants of human β-cell failure remain elusive. Using regulatory-network-based single-cell analysis of human islets, we identify aberrant, diabetes-enriched transitional states characterized by metabolic inflexibility, α/β-transition, and endocrine progenitor/stem cell features. A coordinated transcription factor hierarchy mediating cell state transition emerged and was validated using barcoded guide-based, single-cell gene transfer and calcium flux measures in primary human islet cells. Specifically, two master regulators and associated epigenetic drivers emerged, one (AFF3) controlling β- to α-like-cell reprogramming, the other (BACH2) transition to a dedifferentiated endocrine progenitor-like cell. The findings provide mechanistic insight into diabetic islet cell dysfunction and suggest actionable pathways for pharmacological intervention.

## INTRODUCTION

Type 2 diabetes (T2D) is characterized by defects of insulin action and production^1^. The latter include pancreatic β-cell function and mass abnormalities, which in turn likely impinge on the function of other pancreatic hormones, primarily glucagon^2^. Unveiling mechanisms of islet cell dysfunction is important for our ability to design mechanism-based, durable, and safe disease-modifying interventions. Theories on β-cell failure abound^3^. Among them, dedifferentiation of mature β-cells into endocrine progenitor-like cells^4^, with an attendant increase in cell heterogeneity, has been demonstrated by cell lineage tracing in rodents^5, 6^. Human autopsy studies are consistent with this hypothesis^7–9^ but, in view of morphological and functional differences between the two species, cannot be considered dispositive.

That β-cell undergo dedifferentiation is teleologically attractive, in view of the clinical features associated with β-cell failure: limited adaptive potential of β-cell mass^10, 11^, increased glucagon “tone”^2^, rapid progression in response to incipient hyperglycemia^12^, and prompt reversibility upon treatment, at least in the initial phases of the disease^13^. In this regard, recent clinical trials on diabetes reversal can be reconciled with the idea of β-cell dedifferentiation^14^.

Unfortunately, the limited ability to access the endocrine pancreas in vivo, let alone in a prospective fashion, hinders our ability to functionally interrogate islet cells. Several groups have thus sought to address this problem using computational cell state clustering at the single-cell level^15–21^. Yet, while these studies have demonstrated islet cell gene expression heterogeneity^18, 20, 22^, a clear diabetic islet cell signature remains elusive.

To circumvent these limitations^23^, we employed a systems biology approach to leverage accurate regulatory networks—as inferred by the ARACNe algorithm^24^—for the identification of master regulator (MR) proteins representing mechanistic determinants of gene expression signatures^25^ associated with diabetic vs. non-diabetic islet cell state at the single-cell level. Algorithmic predictions were then experimentally validated in a biologically relevant human primary cell context. This approach, while extensively and successfully tested to elucidate mechanistic determinants of a variety of cancer- and neurodegenerative-related phenotypes^26^, had not yet been applied to the study of metabolic diseases.

Using single-cell RNA-seq (scRNA-seq) from human islets—harvested from either normal (ND) or T2D donors—we first generated islet-cell-specific transcriptional regulatory networks that were then used to measure the transcriptional activity of ∼5600 transcriptional regulator proteins—including transcription factors, co-factors, and signal-transduction proteins— using an extension of the VIPER^25^ algorithm for single-cell analysis (metaVIPER^27^). Basically, the activity of each protein is measured based on the expression of its transcriptional targets, akin to a multiplexed gene-reporter assay.

Protein-activity-based cluster analysis identified multiple, transcriptionally-distinct signatures, representing physiologic, *β−* and *α−*cells related states, as well as aberrant, intermediate transcriptional states that were highly enriched in T2D-derived cells. The latter were characterized by: (*a*) metabolic inflexibility/stress response, (*b*) mixed α/β-cell identity, and (*c*) endocrine progenitor/stem cell features. The analysis also identified MR proteins representing putative mechanistic determinants of the aberrant, T2D-related transitional states. These analytical predictions were validated by gain-of-function studies in primary β-cells, using a single-cell, ectopic gene expression methodology, where TFs were screened for their ability to convert ND into T2D-enriched cells. Finally, to interrogate β-cell function of fate-converted cells, we employed single-cell measurements of intracellular calcium influx in response to glucose. By so doing, we identified TFs drivers of different cellular phenotypes, and obtained a comprehensive functional map of cellular abnormalities in diabetic islets.

## RESULTS

### Heterogeneity of ND and T2D islet cells

We analyzed single cell populations from four non-diabetic controls (ND) and six T2D islet donors of different ages, body mass index (BMI), known duration and control (HbA_1c_) of disease (Supplementary Table 1). ScRNA-seq profiles were generated using the Fluidigm C1 high-throughput integrated fluidic circuits (HT IFC). Since β-cell number is reduced in T2D islets^3^, we enriched samples for β-cells via FAD-mediated autofluorescence-activated cell sorting (Supplementary Fig. 1), and independently analyzed scRNA-seq profiles from either β-cell-enriched samples or whole islets. This approach supported a comprehensive assessment of the β-cell repertoire in T2D patients, while also providing an unbiased assessment of islet cell composition at the single-cell level. After loading cells onto C1 HT IFCs, we visually inspected individual chambers to exclude multiple cells captured into the same chamber. A total of 6,137 cells met quality control criteria and were analyzed further.

T-SNE plotting of raw single-cell mRNA data revealed that individual sample variation and batch effects (color-coded, Fig. 1A) trumped cellular identities, as indicated by the emergence of single-cell clusters that were largely independent of disease status. To buffer donor-to-donor variation and batch effect, we thus relied on the network-based metaVIPER algorithm, which has been shown to be highly effective in removing systematic sample-to-sample bias and batch effects that do not reflect the biology of the cellular state^27^.

**Figure 1.**
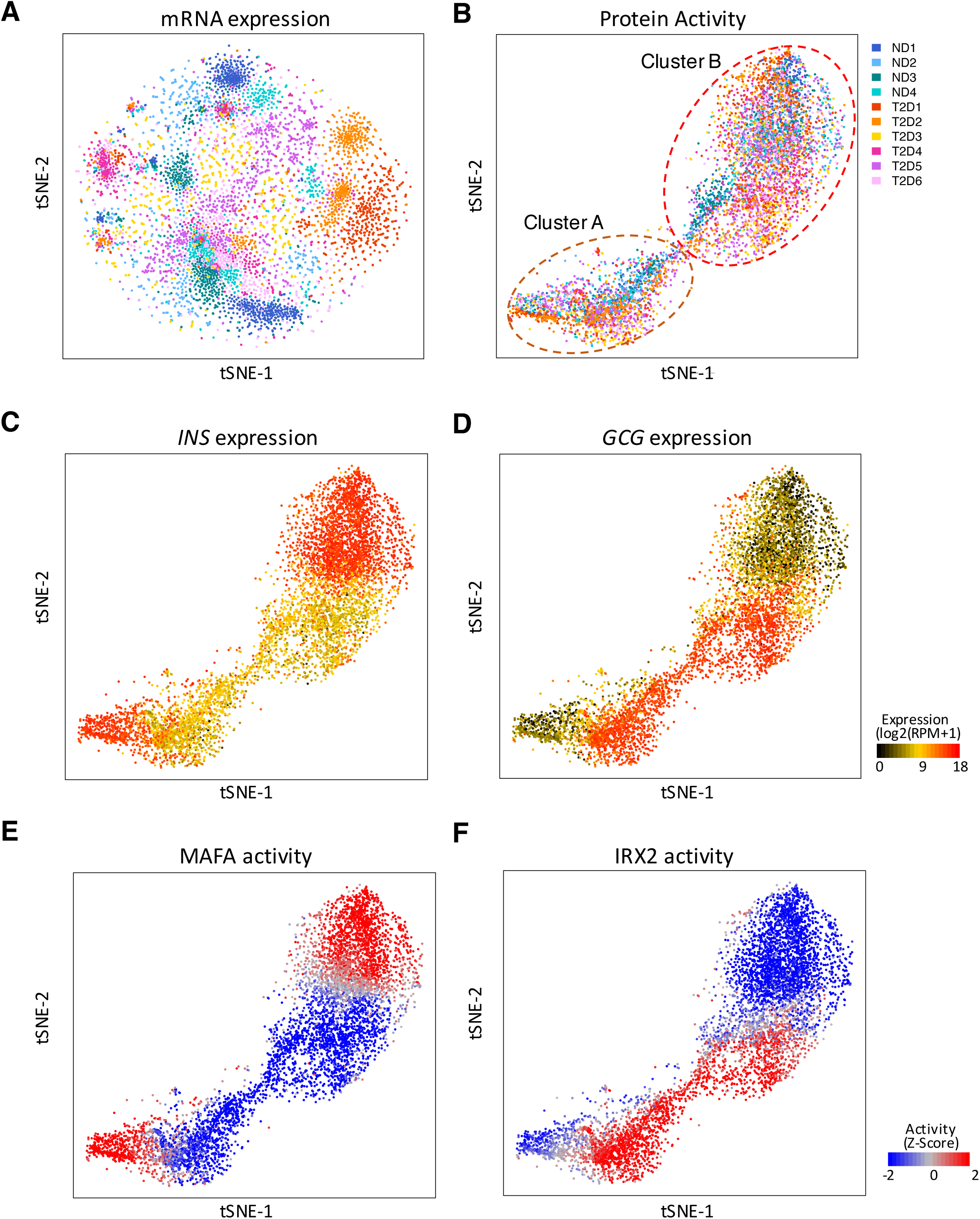
metaVIPER analysis reduces donor-to-donor variability. (A) Single cells from human ND and T2D islets were projected onto 2-D t-SNE space based on whole mRNA expression. Each dot represents a single cell, color-coded according to donor. (B) t-SNE clustering as in (A) but based on protein activity inferred from islet specific regulatory networks. (C) *INSULIN* and (D) *GLUCAGON* mRNA expression plotted in t-SNE at the single cell level. (E) Computationally-inferred MAFA and (F) IRX protein activity plotted in t-SNE at a single-cell level.

Transcription factors (TF) are key to understand cellular identity, but tend to be expressed at much lower levels compared to most cell surface receptors or enzymes in terminally differentiated cells^28^. This poses a challenge when trying to reliably detect subtle differences in TF levels due to the very low sequencing depth of scRNA-seq profiles. Indeed, most genes (in some cases >80%) are not detected by even one transcript (i.e., gene dropout) and most detected genes are supported by a single read^29^. As a result, reliable differential gene expression assessment is challenging. As previously shown^27^, metaVIPER is especially well suited to overcome this limitation because the activity of a protein is measured from tens to hundreds of its transcriptional targets, thus allowing reliable protein activity assessment from profiles with only 20K to 50K reads. Indeed, Spearman correlation between a 30M and a 50K read, VIPER-inferred protein activity profile is ρ = 0.85, while for gene expression it is only ρ = 0.3^25^. In particular, metaVIPER allows quantitative activity assessment of proteins whose encoding mRNA is not detected by even a single read, thus allowing characterization of key lineage marker proteins in virtually every single cell.

These algorithms rely on accurate mappings of proteins’ transcriptional targets (regulons). To address this, we used the ARACNe algorithm^24^ to assemble islet cell-specific regulons using 1,813 transcription factor (TF), 969 cofactor (co-TF), and 3,370 signal transduction (ST) proteins. We then performed metaVIPER analysis, using these regulons, to measure TF/co-TF activity in normal and T2D islet cells. As expected, protein-activity-based t-SNE analyses largely removed the batch effect of individual samples and were effective in identifying clusters associated with two major populations (clusters A and B) (Fig. 2B), without noticeable donor-to-donor variation. Consistent with prior results^27^, this shows that protein-activity-based cluster analyses can buffer technical bias and batch effects typical of scRNA-seq profiles to reveal hidden, biologically-relevant population structures.

**Figure 2.**
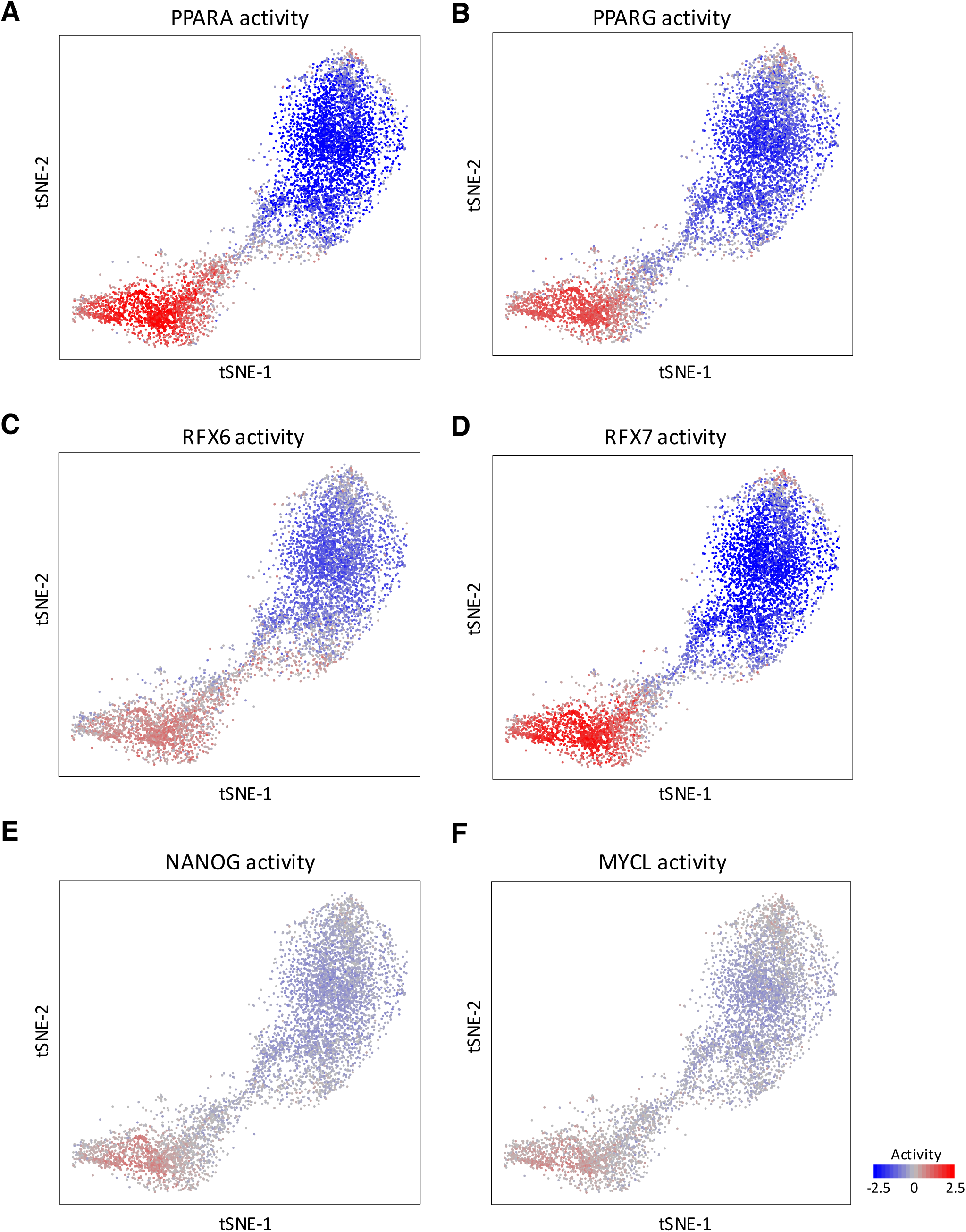
Metabolic inflexibility and stemness markers define two major clusters in ND and T2D islet cells. (A, B) Selected markers of metabolic inflexibility PPARα (A) and PPAR*γ* (B) protein activity plotted in t-SNE. (C, D) Endocrine progenitor marker, RFX6 (C) and its cognate factor RFX7 (D) protein activity presented in t-SNE space. (E, F) Stemness markers, NANOG (E) and MYCL (F) protein activity plotted in t-SNE at a single-cell level.

Clusters A and B were largely independent of disease status, as evidenced by the distribution of color-coded donors (Fig. 1B and Supplementary Fig. 2). Surprisingly, however, they did not recapitulate the β- and α-cell populations of pancreatic islets, as shown by integrative assessment of *INS* and *GCG* mRNA expression levels (Fig. 1C and 1D) and MAFA and IRX2 protein activity. The latter represent well-established markers of β- and α-cells that show mutually exclusive activity in single cells (Fig. 1E and 1F). Indeed, *INS* and *GCG* mRNA matched perfectly with MAFA and IRX2 activity, respectively, showing that both clusters could be further divided into β- and α-cell sub-clusters—thus suggesting a deeper, hierarchical single-cell structure—and confirming metaVIPER’s ability to effectively measure established marker activity (Fig. 1C and 1D). As shown in Supplementary Fig. 3, activity of these proteins would have been virtually undetectable in a majority of cells based on single-cell expression of their encoding genes.

### Biological signatures of islet cell clusters

To assess the biology of clusters A and B (Fig. 1B), we further surveyed their differentially active proteins (Supplementary Data 1). Gene Ontology (GO) analyses of cluster A *vs*. B (illustrated in a REVIGO summary in Supplementary Fig. 4)^24, 25, 27^, shows highly significant enrichment for GO terms such as metabolism, development, and differentiation in cluster A. Among the proteins subsumed under metabolism in GO, we detected a striking activation of PPARα and PPAR*γ* (Fig. 2A and 2B), two markers of metabolic inflexibility–a stage in the progression of β-cell failure (Fig. 2A and Supplementary Fig. 5)^6^. In addition, FOXO1, HIF1α, HSF1 and TP53 were activated in cluster A, consistent with a metabolic stress response (Supplementary Fig. 5)^30^. We also found activation of RFX6 and RFX7 (Fig. 2C and 2D). RFX6 promotes islet cell differentiation, and its mutations are associated with islet agenesis^31^ and diabetes^32^ in humans. Furthermore, TF proteins associated with cell stemness (NANOG, MYCL, and POU5F1) were differentially active in a tightly-clustered subset of RFX6/7-positive cells (Fig. 2E and 2F), especially in T2D islets. Thus, cluster A appears to be comprised of endocrine progenitor-like cells, displaying stem cell features, a prominent signature of β-cell dedifferentiation^4, 33–35^. These results suggest that the two main functional clusters in human ND and T2D islets are defined by drivers of metabolic inflexibility/stress response, and endocrine-progenitor/stem-cell-like features.

### Sub-clustering analysis

To explore the fine-grain substructure of clusters A and B, we performed hierarchical clustering using the iterCluster algorithm^36^. The resulting sub-cluster architecture is visualized as a hierarchical-clustering heatmap (Fig. 3A and Supplementary Fig. 6), and as t-SNE plots (Fig. 3B-3D). To refine functional characterization of individual sub-clusters, we mapped single-cell-specific features, including (*a*) hormone expression (*INS*, *GCG*, and *SST*), as well as activity of TFs associated with (*b*) either β-(MAFA, PDX1, NKX2.2, NEUROD1) or α-cell identity (IRX2, ARX, GLI,3, IGFBP2, ITGB8, HSPB1, F10, SPOCK3, MYO10, CLU)^37^, (*c*) metabolic inflexibility/stress response (PPARα/*γ*, FOXO1, RB1, FOXM1)^6^, and (*d*) endocrine progenitor-(PAX4, PAX6, ISL1, RFX6/7, HES1)^38^ or stem-like cell identity (POUF5F1, MYCL, NANOG).

**Figure 3.**
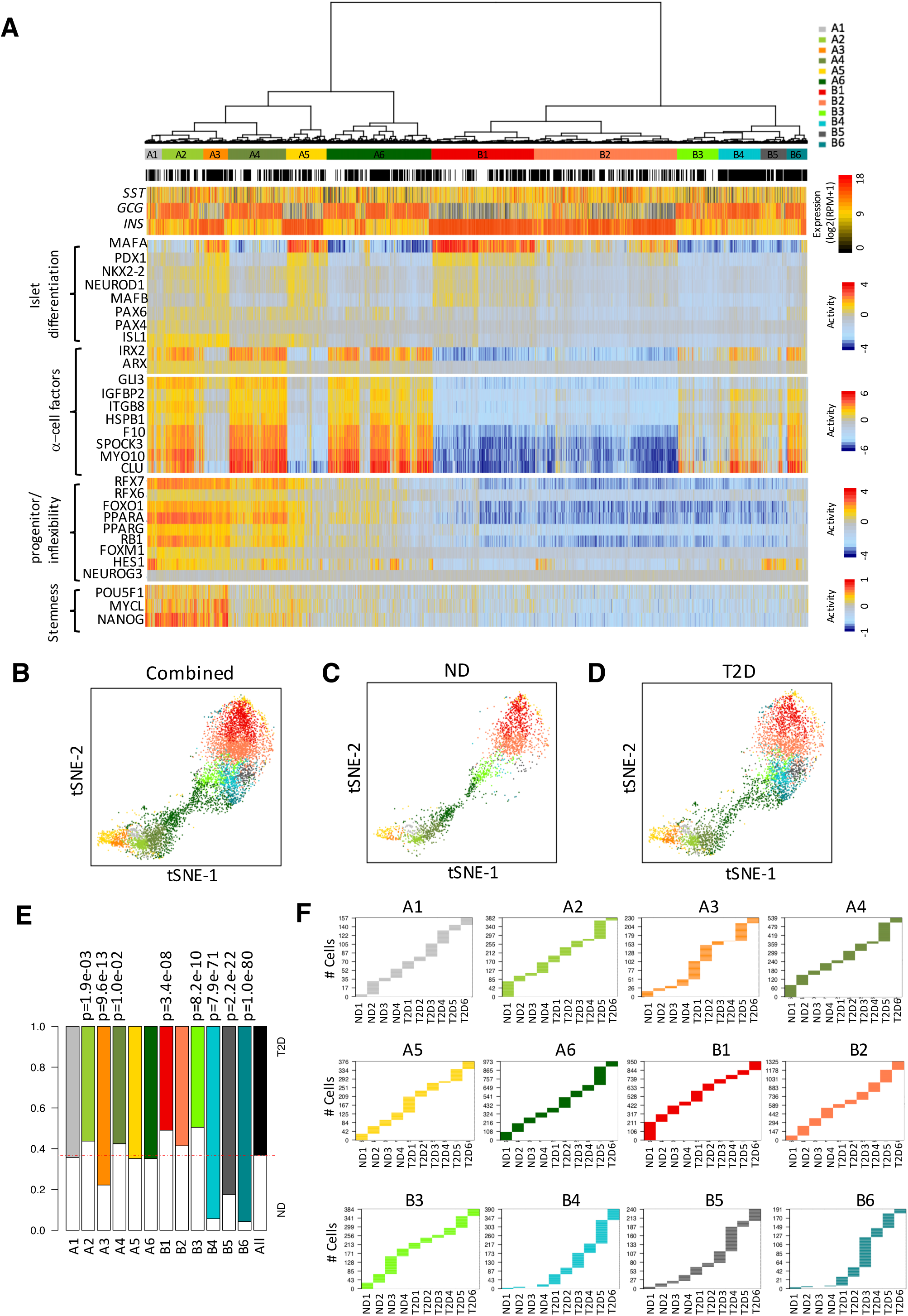
iTerClust classifies ND and T2D islet cells in different biological states. (A) hclustering using the iterCluster algorithm structures cell positioning and sub-groups. Each sub-group is color-coded. Each bar denotes a single cell. Black bars represent T2D cells and white bars ND cells. *SST*, *GCG* and *INS* mRNA expression is plotted at a single-cell level. (B-D) For illustration purpose, sub-clusters were projected onto 2D t-SNE space according to metaVIPER inference as ND and T2D combined (B), or as ND only (C) and T2D only (D). (E) Bar-plots presenting the percentage of ND or T2D cells in each sub-cluster. A dashed line represents the proportion of cells from ND or T2D islets subjected to scRNA-seq analyses. (F) Bar-plots showing the number of cells from individual donors in each subcluster.

This analysis identified six molecularly-distinct sub-clusters of cluster A (A1-A6) and six of cluster B (B1-B6) (Fig. 3A). Four clusters displayed established β-cell features: B1, B2, A3, and A5. Specifically, B1 contained the healthiest β-cells, as indicated by the highest level of *INS* expression and β-cell-specific TF activity, as well as by inactivity of proteins associated with metabolic inflexibility/stress response, as well as progenitor/stem cell and α-cell identity; B2 had similar, yet less pronounced β-cell-specific TF features. In contrast, A3 and A5, while retaining β-cell features, also showed activation of proteins associated with metabolic inflexibility/stress response and endocrine progenitor-like cells, which were more pronounced in A3 than A5. The latter phenotype strikingly recapitulates the response documented in diabetic rodent islets, as discussed below (Fig. 3A)^6^.

In contrast, eight clusters showed clear α-cell-like features: A6 showed the strongest α-cell-like identity, with high *GCG* expression and IRX2 activity, modest activation of proteins associated with metabolic inflexibility/stress response, and virtually no β-cell-like features. While retaining α-cell-like features, A4 and A2 presented strong activation of proteins associated with metabolic inflexibility/stress response and endocrine-progenitor identity and, in cluster A2, TFs associated with stem-cell-like identity. A1 was similar to A2, but showed a mixed pattern of *INS* and *GCG* expression (Fig. 3A). Finally, B3-B6 showed weaker α-cell identity, characterized by *GCG* expression but low IRX2 and metabolic inflexibility/stress response activity. When the two main α-cell-like sub-clusters are compared (A2, A4, A6 *vs*. B3-6), strong IRX2 activity was significantly associated with activation of metabolic inflexibility/stress response proteins (*p* = 1.5E-4, by Chi-square test).

### T2D-enriched sub-clusters

Next, we asked how disease status co-segregated with the 12 molecularly-distinct clusters identified by our analysis. t-SNE plots revealed highly distinct cell distribution patterns between ND and T2D islets, with additional cell sub-populations and cluster representation differences in the latter (Fig. 3C and 3D). We excluded batch effects as potential confounding factors, since all T2D donors share consistent t-SNE patterns that are distinct from t-SNE patterns of individual ND donors (Supplementary Fig. 2C).

We thus proceeded to quantitatively assess differential representation and associated statistical significance of ND- and T2D-derived cells across the 12 clusters (Fig. 3E and 3F). Four clusters were significantly enriched in ND islets: B1, B3, A2, and A4. B1 (*p* = 3.4E-8) corresponds to the healthiest β-cells. B3 (*p* = 8.2E-10) shows an unusual pattern, characterized by both *INS* and *GCG* expression, without activation of either β- or α-like transcriptional programs. We surmise that they are comprised of functionally quiescent cells. A2 (*p* = 1.9E-3) and A4 (*p* = 1.0E-2) also displayed α-like features and activation of the metabolic inflexibility/stress response. A2 shows progenitor/stem-cell-like features. Overall, ND islets display a greater share of healthy β-cells, as well as cells that combine transitional *β/α*, metabolic inflexibility/stress response and progenitor/stem cell-like features. Total cell counts for each cluster in each donor are shown in Fig. 3F.

Four clusters were significantly enriched in T2D islets: A3 and B4-6. A3 (*p* = 9.6E-13) is characterized by β-cell, metabolic inflexibility/stress response, and progenitor/stem-cell-like features. It is interesting to compare this cluster with the “less healthy” β-cell cluster, B2, which doesn’t show disease-state enrichment. In the latter, MAFA activity levels are low but proteins associated with metabolic inflexibility/ stress response are not activated. In contrast, A3 shows higher MAFA activity and activation of metabolic inflexibility/stress response proteins. This is reminiscent of a model in which, in response to declining β-cell function (i.e., lower MAFA activity), FOXO1 is activated to reboot MAFA^30^. However, this also heralds the onset of metabolic inflexibility (PPARα/*γ*)^6^, as well as a drift towards dedifferentiation^4^, as indicated by the activation of RFX6^31^.

Clusters B4-6 (p = 7.9E-71, 2.2E-20, and 1.0E-80) are the most enriched in T2D-related cells, and are rarely found in ND islet cells (Fig. 3E and 3F). Unlike the neighboring, ND-enriched cluster B3, these clusters display significant IRX2 and (in B5 and B6) HES1 activity, an inhibitor of the endocrine differentiation program^3^. These findings suggest that diabetic islets are enriched in cells that combine weak α-like features with a muted stress response. These may correspond to converted α-like cells, as identified by NKX6.1/ALDH1A3 immunohistochemistry of pancreatic islets in diabetic patients^7^.

### Different driver proteins elicit distinct T2D cell state transitions

To identify MR proteins that mechanistically modulate the transcriptional identity of T2D-enriched sub-clusters, we assessed transcriptional regulators whose targets (including activated and suppressed genes) were over- or under-expressed in cluster B6 (the most representative T2D cluster), compared to well-defined β- and α-cells (clusters B1 and A6, respectively). These proteins have the highest metaVIPER-measured differential activity in the two comparisons and are thus inferred as mechanistic determinants of the differential expression signature between these clusters (Supplementary Data 2).

Interestingly, we detected several TFs with significantly increased activity in T2D-enriched clusters but no TFs with decreased activity, suggesting that transition to a diabetic phenotype is associated with gain-of-function (Supplementary Data 2). We selected 10 candidate MRs for experimental validation based on their activity in T2D-enriched clusters and their association with diabetes susceptibility loci in GWAS studies. They included: AFF3^40^, BACH2^41^, BNC2^42^, GAS7^43^, MYT1L^44^, NFATC3^45^, RFX7^46^, TSHZ2^47^, ZRANB3^48^, and ZNF385D^49^. Critically, their mRNA expression was not significantly different based on scRNA-seq analysis, suggesting that these proteins could not have been identified using more traditional approaches (Supplementary Data 3).

To functionally interrogate these candidates, we performed gain-of-function studies in primary human ND islets using a modification of Perturb-seq, a CRISPR-Cas9-mediated scRNA-seq screen^50, 51^, which we termed single-cell gain-of-function sequencing (scGOF-seq) (Fig. 4B). This approach allowed us to evaluate the consequences of the increased activity of a candidate gene at the single-cell level in a physiologically relevant context. We generated bicistronic adenoviral vectors to express candidate genes and a BFP reporter followed by a unique 18-nt barcode with a PolyA signal (Supplementary Fig. 7). The transcribed DNA barcode maps the targeted gene to an individual cell, allowing its identity and the effect of the gene manipulation to be read out by scRNA-seq and analyzed by metaVIPER. To streamline the experiments, we transduced islets with pooled adenoviruses, each pool containing three constructs (See methods). This allowed us to not only simultaneously screen several candidates, but also to test potential epistatic interactions in cells that incorporated more than one barcode.

**Figure 4.**
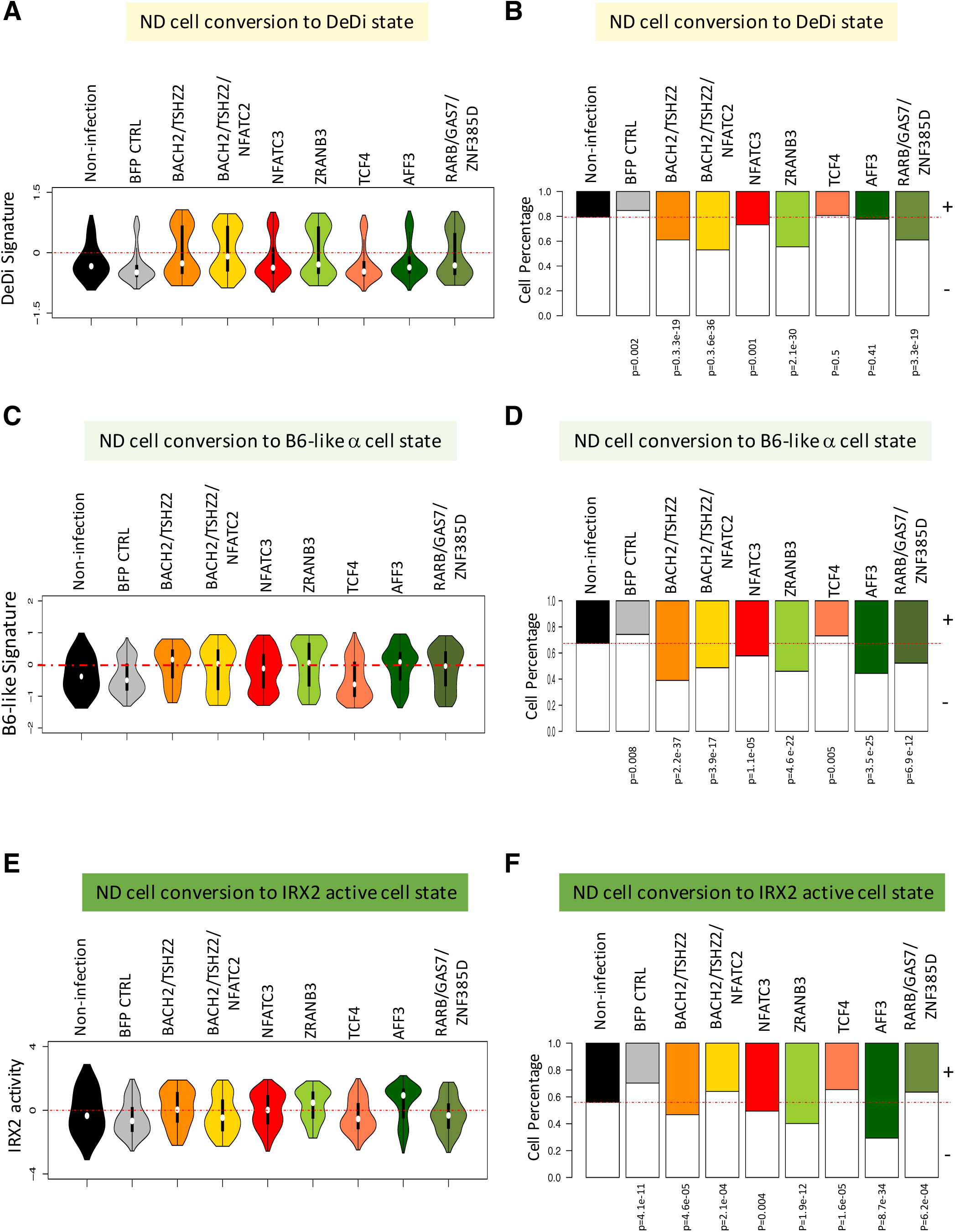
scGOF-seq experiments in human islets. (A) Violin plots showing the distribution of cells following transduction with each individual candidate or combination thereof analyzed according to DeDi signature, an intergrated value of RFX6, RFX7, FOXO1, PPARα, PPAR*γ*, RB1, POUF51, NANOG and MYCL protein activities. Non-transduced and BFP-transduced ND islets serve as negative controls. (B) Bar-plots showing the proportion of islet cells with a positive DeDi signature (> activity 0) in each scGOF-seq condition. A red dashed line indicates the percentage of islets cells with a positive DeDi signature in non-transduced negative controls. (C) Violin plots showing cells with a B6-like signature, which is an intergrated value of IRX2, ZNHIT1, ZFPL1, PAX6 and DRAP1. (F) Bar-plots showing the proportion of islet cells with a B6-like signature as in (B). (E) Violin plots showing cells with IRX2 activity as in (A). (F) Bar-plots showing the proportion of islet cells with a positive IRX2 activity as in (B).

We interrogated candidates for their ability to convert the transcriptional identity of ND islet cells to that of the two most characteristic T2D states: A3 (β-like with metabolic inflexibility/stress response and progenitor/stem-like features) and B6 (putatively converted, α-like) (B6-like signature in Fig. 4). As an integrated measure of the A3 sub-cluster, we used the combined activities of RFX6, RFX7, FOXO1, PPARα, PPAR*γ*, RB1, POUF51, NANOG and MYCL, referred to as DeDi signature. As a measure of B6, we used the combined activities of IRX2, ZNHIT1, ZFPL1, PAX6 and DRAP1, referred to as B6-like signature (Fig. 4). We monitored preservation of β-cell identity throughout the procedure by comparing with non-transduced and BFP-transduced cells (black and gray plots, respectively). The data are presented as violin plots illustrating the shift from the basal (non-transduced or BFP-transduced) to the scGOF-seq cell population, along with quantitative bar graphs (Fig. 4 and Supplementary Fig. 7). In some experiments, the number of cells transduced with individual TFs was too low to allow independent evaluation of each individual factor, due to the stringent cutoff imposed for barcode calling required to remove cells with low candidate-gene expression.

The strongest effects to drive the DeDi signature were seen in cells transduced with the BACH2/TSZH2 or RARB/GAS7/ZNF385D combinations, as well as ZRANB3 (Fig. 4A and 4B and Supplementary Fig. 8). Next, we asked which TFs can drive conversion to α-like-cells (B6-like). AFF3 showed the strongest effect, followed by the BACH2/TSHZ2 combination and ZRANB3 (Fig. 4C and 4D). To understand this latter effect in more detail, we did a sub-analysis asking whether these TFs affected directly the IRX2 network, and found that indeed AFF3 directly induced IRX2 activity (Fig. 4E and 4F and Supplementary Fig. 8). These data suggest that an active transition to an α-like cell state is characteristic of T2D.

To determine the mechanism of the observed fate transition to a DeDi signature, we analyzed TFs whose activity changed upon conversion. We found that BACH2, FOXO1, MYTL1, NFATC3, RFX7 and TCF4 were co-regulated in all conditions conducive to increasing the DeDi signature (Supplementary Fig. 9). While FOXO1 and TCF4 ectopic expression didn’t drive the DeDi signature, their activity significantly increased upon BACH2 or ZRANB3-dependent conversion, suggesting that these six TFs form a hierarchical module driving conversion. This observation is consistent with RNA-seq analyses of β-cell-specific FoxO1 knockout mice, showing impaired levels of *Bach2* mRNA^5, 6^.

In a replication experiment using islets from a different donor, we confirmed that gain-of-function of BACH2/NFATC3/TSHZ2, or CUX2/RFX7 increased DeDi signature cells, as did co-expression of ZRANB3/BNC2/MYT1L (Supplementary Fig. 10).

### Functional effect of master regulators on fate-converted cells

Finally, we tested whether converting ND β-cells to a T2D-enriched signature impairs β-cell function, as assessed by measuring intracellular Ca^2+^ flux in response to glucose at the single-cell level. We selected BACH2 (due to its link to diabetes susceptibility^52^), AFF3 (due to the strong effect on α-like-cell conversion), and TCF4 (as a negative control) for these experiments. To gate β-cells, we co-transduced ND islets with RIP-zsGreen together with adenovirus expressing each candidate TF^53^. After three days, we partially dissociated and plated islets for Ca^2+^ imaging. We individually gated zsGreen/BFP-double-positive cells to monitor intracellular Ca^2+^ concentration based on the Ca^2+^ indicator, Rhod-2 signal (Fig. 5A). β-cells transduced with Tag-BFP showed a brisk Ca^2+^ response to glucose or KCl (Fig. 5B). The glucose response was blunted in cells transduced with AFF3 or BACH2, while the KCl response was intact, indicating that AFF3 or BACH2 gain-of-function impairs β-cell glucose sensing but does not alter membrane depolarization (Fig. 5C and D). This is consistent with the loss of β-cell features observed in scGOF-seq experiments with AFF3 or BACH2 (Fig. 4D). In contrast, TCF4 gain-of-function had no statistical effect on Ca^2+^ influx compared to Tag-BFP (Fig. 5E), consistent with the scGOF-seq results, showing that TCF4 failed to affect β-cell identity. These data further validate our approach to functional testing and strengthen the observations on AFF3 and BACH2.

**Figure 5.**
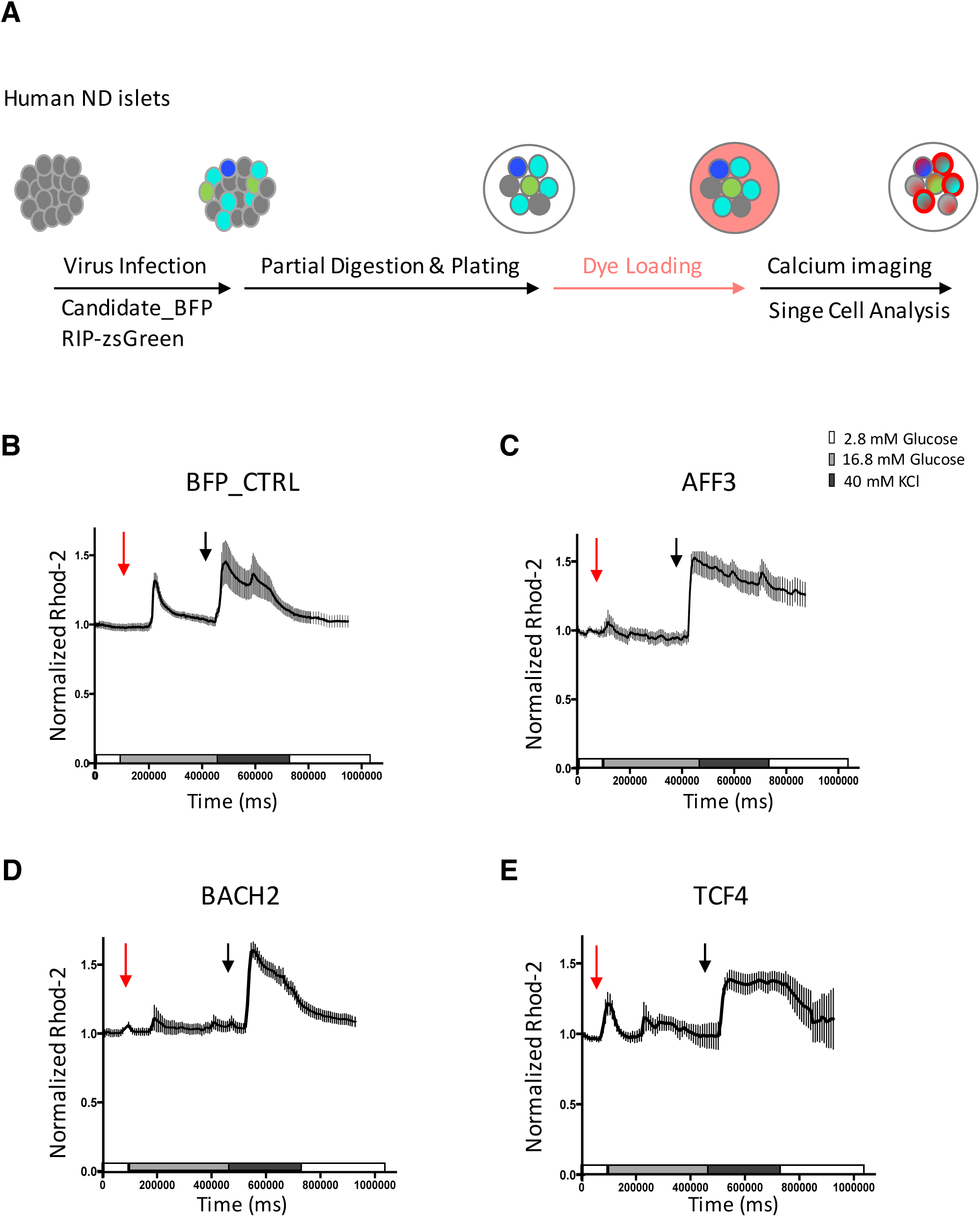
Single-β-cell Calcium microfluorimetry. (A) Schematic drawing of the single-cell Ca^2+^ imaging procedure. (B)-(E) Representative traces of Ca^2+^ flux measured by Rhod2 loading in Ad-BFP (B), Ad-AFF3 (C), Ad-BACH2 or Ad-TCF4 transduced primary human *b*-cells. Red arrows indicate the timing of addition of 16.8mM glucose, and black arrows indicate addition of 40mM KCl.

## DISCUSSION

The clinical progression of T2D is characterized by a rapid degradation of β-cell function^12^ that becomes refractory to treatment over time, requiring combination therapy and resulting in life-threatening complications. Glp1-based agents have β-cell protective effects, but don’t appear to significantly delay monotherapy failure, or reverse established β-cell failure^54^. Our work aims to identify mechanistic networks capable to reverse β-cell dedifferentiation as a disease-modifying approach^14^.

Unbiased analysis of cell transition states identified three salient features of islet cell populations: (*a*) metabolic inflexibility/stress response (FOXO1 and PPARα/*γ*); (*b*) endocrine progenitor (RFX6/7) and stem-cell-like features (NANOG, L-MYC expression); and (*c*) α-cell-like features (*GCG* expression, activation of IRX2 and related regulators). Within this distribution, healthy β-cells (sub-cluster B1) are enriched in ND subjects, while β-cells with activated metabolic inflexibility/stress response, as well as progenitor/stem cell features are enriched in T2D (sub-cluster A3). In the latter sub-cluster, activation of the two nuclear hormone receptors PPARα and *γ* may appear counterintuitive, since they oversee different lipid metabolic functions, but is in fact consistent with the observation that, as islets fail, synthetic and oxidative branches of lipid metabolism are coactivated^6^. These data provide a potential explanation for the longstanding clinical observation that PPAR*γ* agonists have beneficial effects on diabetes prevention and β-cell function^55^. While the latter have traditionally been ascribed to improved insulin sensitivity, the present data are suggestive of direct effects on β-cells.

A striking feature of the MR analysis is the existence of intermediate states providing continuity between the main β- and α-cell identities. This can be reconciled with the observation that the latter can be reprogrammed to serve as surrogate β-cells^56^, as well as with the overall plasticity of endocrine islet cell fate. It is striking that T2D islets are enriched in “stressed” β-cells (i.e., β-cells with markers of metabolic inflexibility/stress response/progenitor/stemness), whereas ND islets are enriched in “stressed” α-like-cells. In this regard, there appears to be a direct correlation between *GCG* mRNA, activation of the IRX2 network, and metabolic inflexibility/stress response. Cells in which these features are more pronounced prevail in ND islets, whereas cells with more muted α-like characteristics are enriched in T2D, somewhat reminiscent of mixed-feature cells identified previously^7^. What can this mean? One potential interpretation is that α-like cells displaying a more robust stress response are more resilient, hence capable of reversal to β-cells (if indeed this is their origin), whereas those lacking a stress response are transitioning toward a functionally quiescent state.

The biological relevance of the hierarchical clustering is supported by functional studies in primary islets, showing that imputed regulatory networks can drive different aspects of the cellular pathology. In this regard, our data favor a network model in which a small subset of TFs can drive different aspects of the diabetic islet pathology. Thus, BACH2/TSHZ2 strongly affect dedifferentiation, whereas AFF3 has the strongest effect on driving the α-like-cell phenotype. This analysis is strengthened by the functional data on calcium fluxes, indicating that gain-of-function of BACH2 or AFF3 does indeed impair the features of a differentiated β-cell.

More is known about the function of BACH2 than that of AFF3 in islets. Traditionally viewed as a repressor of lymphocytes lineages^57^, BACH2 has been linked to genetic susceptibility to type 1 diabetes^52^, and changes to its chromatin structure have been reported in human diabetic islets^58^. In addition, it’s regulated by the Akt/mTOR pathway, has been shown to dimerize with MAF proteins^57^, and has emerged as a candidate from a previous analysis of β-cell dedifferentiation^5^, a role confirmed by the present findings. AFF3 has been known to serve as a scaffold for the transcription elongation factor (P-TEFb)-containing super elongation complex (SEC), which regulates rapid gene activation upon exposure to environmental stimuli^59^. It has also been linked to enhancer activation and expression of long noncoding RNA and miRNA genes from the active allele^60^, as well as to neural development^61^.

Previous studies using scRNA-seq have been only partly successful to characterize distinct cell sub-groups in T2D^15–21^, potentially due to low sample size and sequencing depth, as well as untested functional consequences of altered gene expression profiles among different cell sub-populations. In this work, we leveraged the scRNA-seq procedure in two ways. First, we generated and interrogated islet-specific transcriptome regulatory networks using ARACNe and metaVIPER. Moreover, we experimentally validated the systems-level approach by a contextual screen in β-cells. As next steps, we plan to test whether T2D-enriched sub-populations can be reconverted to healthy *β−*cells, and use the OncoTreat method to identify chemical modulators of specific MR activity signatures^62^. Ultimately, this work will expand our knowledge of T2D pathophysiology and pave the way to the identification of treatments to benefit T2D patients.

## METHODS

### Human islet cell culture

Human ND and T2D islets were obtained from the National Institutes of Health’s Integrated Islet Distribution Program (IIDP). Upon arrival, human islets were plated at a density of 10,000 IEQ per 10-cm non-treated tissue culture dish (Corning, Corning, NY; cat. no. 430591) into 10 ml of islet culture medium (Prodo Labs, PIM(S)®, cat#PIM-CS001GMP), supplemented with 5 ml PIM(G)® Glutamine/Glutathione (Prodo Labs, cat#PIM-G001GMP), and 5% PIM(ABS)® Human AB Serum (Prodo Labs, cat#PIM-ABS001GMP), along with triple antibiotics, PIM(3X) ®, which includes Ciprofloxacin (Ref 61-277-RF, 10mg/1000ml), Gentamycin (Sigma, G1272, 10mg/1000ml), and Amphotericin B (Omega, FG-70, 2500mcgm/1000ml). Islets were cultured for no longer than one week after arrival and medium was replaced every 2 days.

### Fluorescence-Activated β-Cell Sorting

On the day of scRNA sequencing, islets were collected, washed in phosphate-buffered saline (PBS) once, and dispersed into single cells by mechanical shaking at 37°C using 0.05% trypsin (Gibco, 25300054). Dispersed islet cells were incubated for 1 hr. at 37°C in MEM (Gibco, 11090081) containing 1% BSA, then collected into 5 mL round-bottom polystyrene tubes with a cell strainer (BD Falcon, 352235) at a final SYTOX Red concentration 5 nM (ThermoFisher, S34859). High- and low-fluorescence cells were sorted on a fluorescence-activated Influx cell sorter (BD Influx system). Flavin adenine dinucleotide (FAD) content of cells was analyzed at an excitation wave length of 488 nm, and collected at 525 nm. For scGOF-seq, adenovirus-transduced cells were further gated for PacBlue-positive cells at an excitation wave length of 401 nm, and collected at 452 nm.

### RNA-seq library preparation using the Fluidigm C1 800 platform

Sorted cells with high or low/medium FAD content were suspended in C1 Cell Suspension Reagent (Fluidigm) and loaded onto each inlet of the C1 high-throughput integrated fluidic circuit (HT IFC). The number of cells captured in each chamber was visualized and noted using a phase contrast microscope. Only chambers with single-cell capture were used for analysis. Cells were lysed, and mRNA reverse transcribed and PCR-amplified using C1 Single-cell Auto Prep IFC (Fluidigm, protocol 101–4964). The quality and yield of cDNA were determined by Agilent Bioanalyzer using Agilent High Sensitivity DNA Chip. Libraries for sequencing were prepared using Nextera XT DNA library preparation kit (Illumina FC-131-1096) and sequenced with paired-end 50 cycles on Illumina HiSeq2500. Each library pool was subjected to 2 lanes of Illumina HiSeq2500.

### 10x Genomics platform RNA-seq library preparation

Sorted cells were treated with a Chromium Single Cell 3’ Library and Gel Bead Kit V2 (PN-120237), Chromium Single Cell 3’ Chip Kit V2 (PN-120236) and Chromium i7 Multiplex Kit (PN-120262) and analyzed with a 10x Genomics Chromium for Single-Cell Library Preparation Instrument, per the manufacturer’s specifications and sequenced paired-end 150 bp on HiSeq 4,000 to a depth of 90,000 UMI per cell. UMI counts for each cellular barcode were quantified and used to estimate the number of cells successfully captured and sequenced. The Cell Ranger Single-Cell Software suite was used for demultiplexing, barcode processing, alignment, and initial clustering of the raw scRNA-seq profiles.

We used the Chromium instrument and the Single Cell 3’ Reagent kit (V1) to prepare individually barcoded single-cell RNA-Seq libraries following the manufacturer’s protocol (10X Genomics). For QC and to quantify the library concentration we used both the BioAnalyzer (Agilent BioAnalyzer High Sensitivity Kit) and qPCR (Kapa Quantification kit for Illumina Libraries). Sequencing with dual indexing was conducted on an Illumina NextSeq machine using the 150 cycles High Output kit. Sample demultiplexing, barcode processing, and single-cell 3’ gene counting was performed with the Cell Ranger Single Cell Software Suite CR2.0.1. Each droplet partition’s contents were tagged with a unique molecule identifier (UMI) – a barcode encoded as the 2nd read of each sequenced read-pair. We followed the 10X Single Cell 3’ Reagent Kits v2 protocol as written, using 12 cycles for cDNA amplification and 12 cycles for sample index PCR. Samples were sequenced to a depth of ∼400M reads per sample on a NovaSeq 6000 (R1 = 26bp, R(i) = 8bp, R2 = 91bp).

### Plasmids

We synthesized open reading frames of each scGOF-seq candidate with a Tag-BFP and an 18-nt unique barcode (Supplemental Table 2) (Qinglan Biotech) and cloned them into the pENTR2b vector using KpnI and EcoRV (AFF3, CUX2, FOXO1, GAS7, TSHZ2 and ZFN385D), BamHI and NotI (BACH2, BNC2, EBF1, RARB, RFX7 and TCF4), SalI and NotI (MYT1L and NFATC3), or KasI and NotI (ZRANB3).

### Adenovirus generation

Recombinant adenoviruses were generated using the pAd/CMV/V5-DEST Gateway recombination system (Life Technologies) after cloning the full-length cDNA into the pENTR vector. Individual adenoviruses were packaged and amplified in HEK-293A cells, then pools of three P1 adenoviruses as detailed below were expanded into high-titer virus. Pool1: ZNF385D, RARB, GAS7; Pool2: EBF1, FOXO1, TCF4; Pool3: BACH2, TSHZ2, NFATC3; Pool4: ZRANB3, BNYC2, MYT1L, and Pool5: AFF3, RFX7, CUX2. Adenoviruses were purified by PD-10 column (17085101, GE Healthcare). Titers were determined by plaque assay (PFU). Each virus pool was transduced into HCT116 cells and expression was analyzed by qPCR. FOXO1, TCF4, NFACT3, RFX7 and AFF3 were amplified individually from P1 adenovirus.

### Adenovirus transduction

200-300 human ND islets were handpicked for each condition of transduction and placed on 5 mL round-bottom polystyrene test tube. Thereafter, islets were washed and incubated with 100 ul serum-free islet culture medium containing 1mM EGTA, and transduced at an MOI 20. After 5- to 6-hr transduction, 1ml of complete islet culture medium with 5% PIM(ABS)® Human AB Serum (Prodo Labs, cat#PIM-ABS001GMP) was added overnight. Islets were then transferred to 60 mm non-treated tissue culture dishes (Fisher Scientific FB0875713A), and medium was replaced with fresh islet culture medium every 2 days for seven days for scGOF-seq experiments, and three days for single-cell intracellular calcium microfluimetry.

### Single-Cell Intracellular Calcium Microfluimetry

Similar-size human islets from non-diabetic donors were handpicked and transduced with adenovirus expressing each candidate cDNA. The day after transduction, islets were partially dispersed using 0.05% trypsin for 5 min at 37°C, then plated on 35mm glass bottom dishes with 10mm microwells (In vitro Scientific D35-10-0-N) pre-coated with fibronectin (Sigma, F1141). Cells were washed with islet media and allowed to rest for two additional days. On the third day, each plate was washed with KRBH buffer and incubated with 2.8 Mm glucose containing KRBH buffer for 30 min, then loaded in the dark with 5 *μ*M Rhod-2, AM (Thermo Fisher R1244) in KRBH buffer. Cells were washed and transferred into a perifusion chamber placed in the light path of a Zeiss Axiovert fluorescence microscope (Zeiss, USA), and perifused with low glucose (2.8 mM), high glucose (16.8 mM), or KCl (40 mM) in KRBH buffer. β-cells were excited by a Lambda DG-4 150 Watt xenon light source (Sutter, Novato, USA), using alternating wavelengths of 340 and 380 nm at 0.5 s intervals, and imaged at 510 nm. For each data set, regions of interest corresponding to the locations of 10-20 individual cells were selected and images were recorded using an AxioCam camera controlled by Stallion SB.4.1.0 PC software (Intelligent Imaging Innovations, USA). Single-cell intracellular Ca2+ mobilization data are plotted as a function of time.

### Quantitative analyses of single-cell gene expression

For single-cell RNA-Seq using Fluidigm C1 system, after demultiplexing, the resulting raw reads were aligned to hg19 reference index by Bowtie2-2.2.6. Aligned reads were sorted and indexed by samtools-1.2. Counts matrices were quantified with R package GenomicFeatures-1.24.5GenomicAlignments-1.8.4 and TxDb.Hsapiens. UCSC. hg19.knownGene-3.2.2 from Bioconductor. Counts matrices were then converted to log2(reads per million reads + 1) matrices as the final gene expression quantification^27^.

For ScGOF-seq using 10x Genomics Chromium, UMI (Unique Molecular Identifier, barcodes each mRNA transcript) matrices were quantified using scPLATE-Seq pipeline, using parameters –umiLen 10 and –barcodeLen 16 considering the UMI and cell barcode length of Chromium, with customized STAR 2.5.1a reference index with parameter --genomeSAindexNbases 7 considering the length of cDNA and sgRNA sequences, including regular UCSC hg38 reference index and cDNA+sgRNA reference index (one cDNA/sgRNA sequence per artificial chromosome). UMI matrices were then converted to log2(transcripts per million transcripts + 1) matrices as the final gene expression quantification^27^.

### Regulatory networks and transcriptional regulator activity analysis

#### Islet-specific regulatory network generation

Donor-specific regulatory networks were reverse engineered by ARACNe^24^. ARACNe was run with 200 bootstrap iterations using 1,813 transcription factors (genes annotated in Gene Ontology molecular function database, as GO:0003700, ‘transcription factor activity’, or as GO:0003677, ‘DNA binding’, and GO:0030528, ‘transcription regulator activity’, or as GO:00034677 and GO: 0045449, ‘regulation of transcription’), 969 transcriptional cofactors (a manually curated list, not overlapping with the transcription factor list, built upon genes annotated as GO:0003712, ‘transcription cofactor activity’, or GO:0030528 or GO:0045449) and 3,370 signaling pathway related genes (annotated in GO Biological Process database as GO:0007165 ‘signal transduction’ and in GO cellular component database as GO:0005622, ‘intracellular’, or GO:0005886, ‘plasma membrane’). Parameters were set to 0 DPI (Data Processing Inequality) tolerance and MI (Mutual Information) p-value (using MI computed by permuting the original dataset as null model) threshold of 10-8. Transcriptional regulator activity profiles were inferred from combined expression matrix (from all 11 donor-specific matrices) by integrating all 11 donor-specific regulatory networks with the metaVIPER algorithm^27^.

#### Dimension reduction, clustering and pseudo-lineage analysis

Dimension reduction was done using both gene expression and metaVIPER-inferred activity profiles. For gene expression, cells were first projected to principal component space using PCA (Principal Component Analysis), further projected to 2-D t-SNE space. T-SNE function in CRAN R package t-SNE-0.1.3 using 1- r(cell-wise Pearson on principal component space) as distance matrix. For metaVIPER-inferred activity, cells were projected to 2-D t-SNE space. T-SNE function in CRAN R package t-SNE-0.1.3 using cell-wise activity dissimilarity as distance matrix^25^.

Clustering analysis was done using iterClust iterative clustering analysis framework^36^. iterClust function in Bioconductor R package iterClust-1.0.2 at metaVIPER-inferred activity level. In total three iterations were done, separating cell populations at different metabolic stress states and cell types sequentially. Using scRNA-seq data from cells with more than 0.1 million mapped reads, we first projected single cells onto on 2D-space with t-SNE.

## Data availability

Single-cell RNA-Seq data using Fluidigm C1 system for all donors have been deposited at the Gene Expression Omnibus (GEO) under accession number GSE98887. ScGOF-seq data using 10x Genomics Chromium platform have been submitted at the GEO.

## Author contributions

J.S and H.D. designed and performed experiments, analyzed data, and wrote the manuscript. A.C and D.A. designed experiments, oversaw research, and wrote the manuscript.

## ACKNOWLEDGEMENTS

We thank the Genome Technology Center at NYU and Sulzberger Genome Center at Columbia for help with scRNA-Sequencing, the Human Islet and Adenovirus Core of the AECOM/MSSM DRC for help in generating the adenoviruses. FACS experiments were performed in the DRC Flow Cytometry Core (S10OD020056). The content is solely the responsibility of the authors and does not necessarily represent the official views of the National Institutes of Health. This work was supported by NIH grants DK64819, DK63608 (Columbia University Diabetes Research Center). J.S. was supported by Kirschstein-NRSA postdoctoral fellowship F32DK117574. We are grateful to members of the Accili and Califano laboratories for insightful discussions of the data. We thank Travis Morgenstern and Dr. Henry Colecraft (Columbia University) for training and equipment for calcium flux imaging. We also thank to Drs. Andrew Stewart, Adolfo Garcia-Ocaña and Peng Wang for providing RIP-zsGreen adenovirus.

## Supplementary Figure Legends

**Supplementary Figure 1. Islet cells sorted by FAD content-based autofluorescence.** β-cells are enriched by sorting for increased auto-fluorescence. In non-diabetic samples (ND, left), ∼40% islets cells are putative β-cells, whereas in diabetic human islets (T2D, right) only ∼20 % are putative β-cells.

**Supplementary Figure 2. T-SNE Density Clustering of human islet cells.** T-SNE Density Clustering of human non-diabetic (A) or T2D islets (B) based on metaVIPER-inferred transcriptional regulator activity profiles. (C) Single cells from individual donors were projected onto 2-D t-SNE space based on metaVIPER-inferred transcriptional regulator activity profiles.

**Supplementary Figure 3. mRNA expression pattern in tSNE plots.** mRNA expression of MafA (A), Irx2 (B), Rfx6 (C), Rfx7 (D), PPARα (E) or PPAR*γ* (F) was visualized as t-SNE plots.

**Supplementary Figure 4. Functional enrichment of GO terms**. Data were analyzed with GO term finder using the list of proteins significantly activated in cluster A compared to cluster B in Figure 1. Functional enrichment was summarized using REVIGO. Enriched terms remaining after the redundancy reduction are represented as scatterplots in a two-dimensional space, which summarizes GO terms’ semantic similarities.

**Supplementary Figure 5. Characterization of cells with metabolic inflexibility/stress response and stem-like cell features.** MetaVIPER-inferred activity of metabolic inflexibility regulators (A) RB1 and (B) FOXO1, stemness markers (C) FOXM1 and (D) POU5F1 plotted as t-SNE. Metabolic stress-related transcriptional regulators, including (E) hypoxic stress-related HIF1A, (F) oxidative stress-related TP53, (G) JNK family, (H) NFKB1, (I) HSF1, and (J) ER stress-related XBP1 plotted as t-SNE.

**Supplementary Figure 6. iTerClust classifies human ND or T2D islet cells in different biological states.** (A) hclustering using the iterCluster algorithm structures cell positioning and sub-groups of islet cells only from ND donors. Each sub-group was color-coded accordingly. *SST*, *GCG* and *INS* mRNA expression is plotted at a single-cell level. (B) Same as (A) but from T2D donors. (C) Sub-clusters were projected onto 2D t-SNE space according to metaVIPER-inference for ND islets. Gray dots were a overlay of T2D cells (D) Same as (C) but for T2D islets.

**Supplementary Figure 7. Heatmap to denote cell identity after scGOF-seq.** (A) Schematic drawing of scGOF-seq plasmids. (B) Cell annotation of scGOF-seq is presented for each group, color-coded for single candidate or combinatorial candidate transduction.

**Supplementary Figure 8. scGOF-seq analyses using ND islets.** (A) Bar-plots showing the proportion of islet cells with positive MAFA activity in each condition. A red dashed line indicates the percentage of islets cells with positive MAFA activity in non-transduced negative controls. (B) Same as (A) but for IRX2 activity. (C) Bar-plots showing the proportion of islet cells with a positive DeDi signature (> activity 0) in each condition. (D) Bar-plots showing the proportion of islet cells with a positive B6-like signature (> activity 0) in each condition.

**Supplementary Figure 9. Core driver network of ND cell conversion into DeDi signature cells.** Violin plots of core driver network, BACH2, NFATC3, MYT1L and TCF4 for each single candidate or in combination according to DeDiSig score.

**Supplementary Figure 10. Biological scGOF-seq replicate.** (A) Bar-plots showing the proportion of islet cells with positive MAFA activity in each gain-of-function condition. A red dashed line indicates the percentage of islets cells with positive MAFA activity in non-transduced negative control. (B) Same as (A) but for IRX2 activity. (C) Bar-plots showing the proportion of islet cells with a positive DeDi signature (> activity 0) in each condition.

**Supplementary Data 1**. List of differentially activated proteins between Cluster A and Cluster B.

**Supplementary Data 2**. (A) List of differentially activated TFs between Cluster B1 vs. A3, or A6 vs. A3. (B) List of differentially activated TFs between Cluster B1 vs. B6, or A6 vs. B6.

**Supplementary Data 3**. List of differentially expressed genes between Cluster B1 vs. A3, A6 vs. A3, B1 vs. B6, or A6 vs. B6.

