## Supplementary Figures for "AFF3 and BACH2 are master regulators of metabolic inflexibility, β/α-cell transition, and dedifferentiation in type 2 diabetes"

**Supplementary Table 1. Donor information**

| Donor | Sex | Age | BMI | HbA1C (%) | Treatment | Duration (Yrs) |
|---|---|---|---|---|---|---|
| ND1 | M | 25 | 26 | N/A | | |
| ND2 | F | 51 | 20 | N/A | | |
| ND3 | F | 53 | 34 | 5.3 | | |
| ND4 | F | 36 | 24 | 5.0 | | |
| T2D1 | M | 59 | 32 | 9.6 | No Tx | 2 |
| T2D2 | M | 58 | 39 | 8.9 | Metformin | 8 |
| T2D3 | M | 51 | 25 | 6.9 | N/A | upon admissoion |
| T2D4 | F | 42 | 28 | 6.7 | Insulin | 6 |
| T2D5 | M | 48 | 44 | 6.6 | N/A | upon admissoion |
| T2D6 | M | 59 | 33 | 6.6 | Metformin | 2 |

**Supplementary Table 1.** Human islet donor information with age, sex, BMI, HbA1C and medical history.

**Supplementary Table 2. scGOF-seq barcode dictionary**

| Gene | Guide Barcode |
|---|---|
| AFF3 | GATTCTGCCACTACTTCG |
| BACH2 | AAGTAGCGCCTAGACGCA |
| BNC2 | TAGAACATCAATCCGGTT |
| CUX2 | TTGCACCGGAAAGTCTGC |
| EBF1 | TACGTGTCCGTATGACAT |
| FOXO1 | CGCAAGTGTAGCATCAGA |
| GAS7 | CTGACCAACCGCAGAAGT |
| MYT1L | AGGACCACTGGACATCCA |
| NFATC3 | GTTGAATTGTGGAGTTAT |
| RARB | TAATCCGTACAGGTGTCA |
| RFX7 | GAGTGCTTAATGTACCCA |
| TCF4 | TGACACGTATTTCGGAGG |
| TSHZ2 | AAATGACCAACTTGACGT |
| ZNF385D | AGCTAGGCCATTTGTATC |
| ZRANB3 | CTAAACTCATAACATAGA |
| tagBFP | CTCCGGTTGCAGAGGCTA |

**Supplementary Table 2.** List of TFs tested in scGOF-seq experiments and corresponding guide barcode for cell annotation.

SupFig. 1

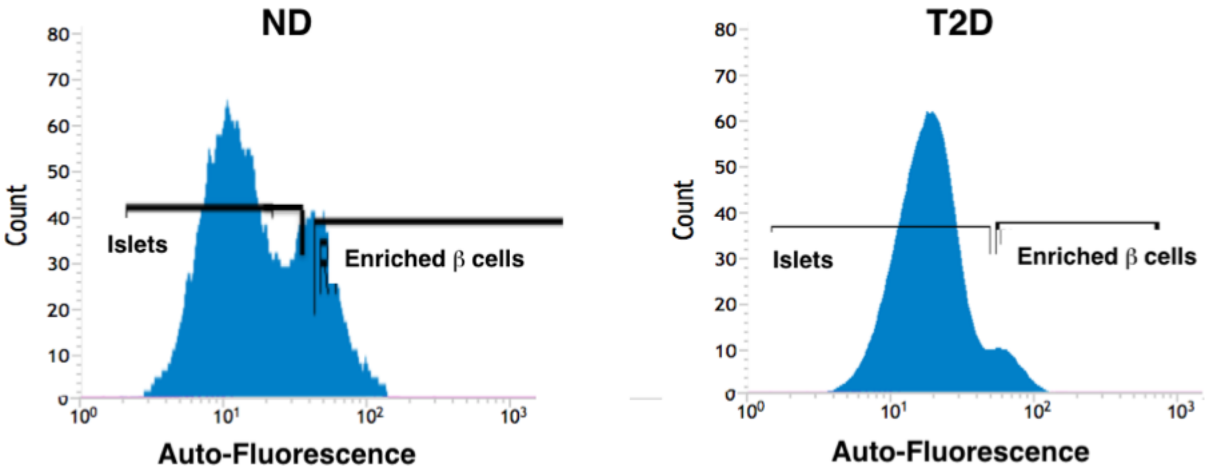


**Supplementary Figure 1. Islet cells sorted by FAD content-based autofluorescence.**
β- cells are enriched by sorting for increased auto-fluorescence. In non-diabetic samples (ND, left), ~40% islets cells are putative β-cells, whereas in diabetic human islets (T2D, right) only ~20 % are putative β-cells.

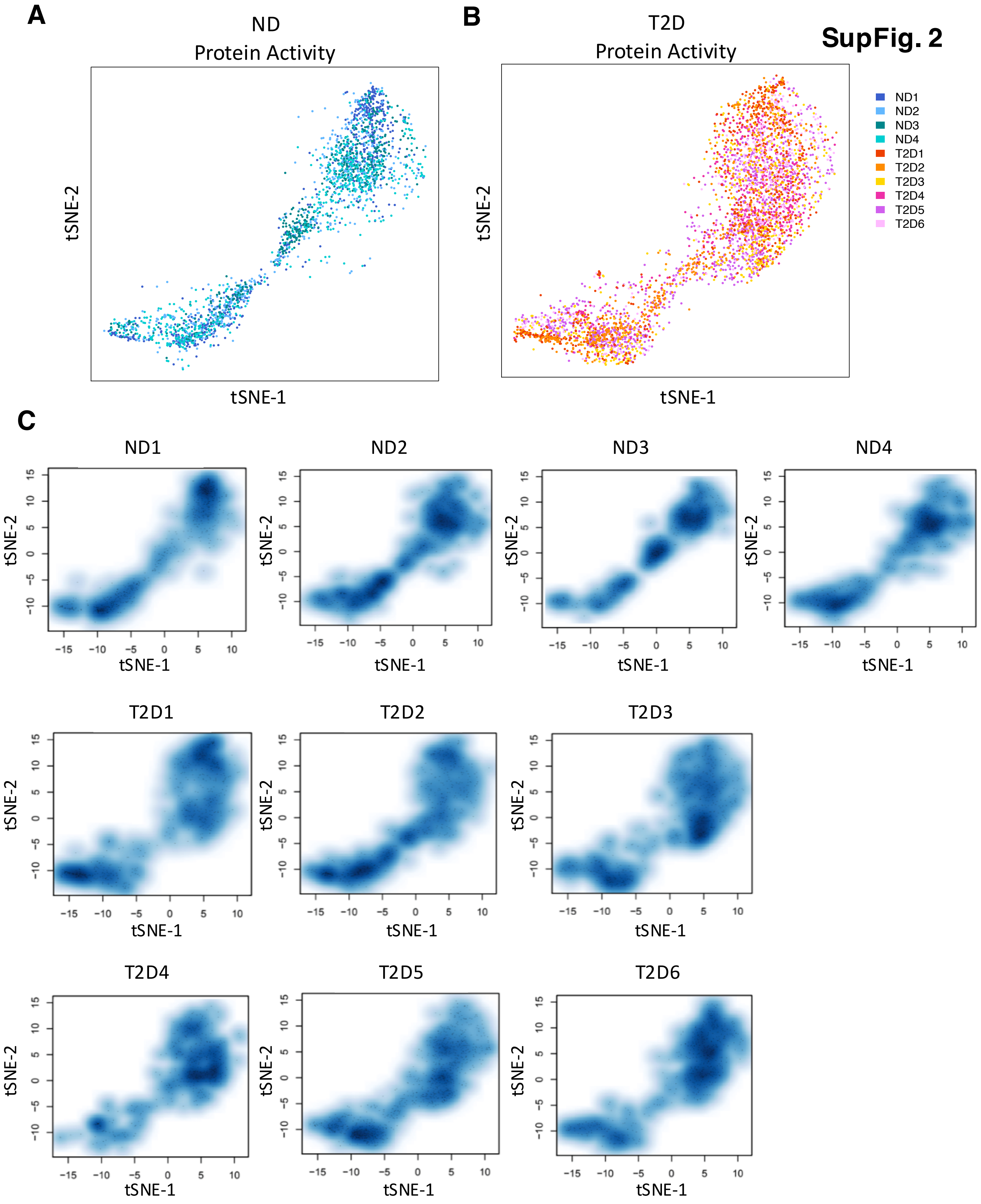


**Supplementary Figure 2. T-SNE Density Clustering of human islet cells.** T-SNE Density Clustering of human non-diabetic (A) or T2D islets (B) based on metaVIPER-inferred transcriptional regulator activity profiles. (C) Single cells from individual donors were projected onto 2-D t-SNE space based on metaVIPER-inferred transcriptional regulator activity profiles.

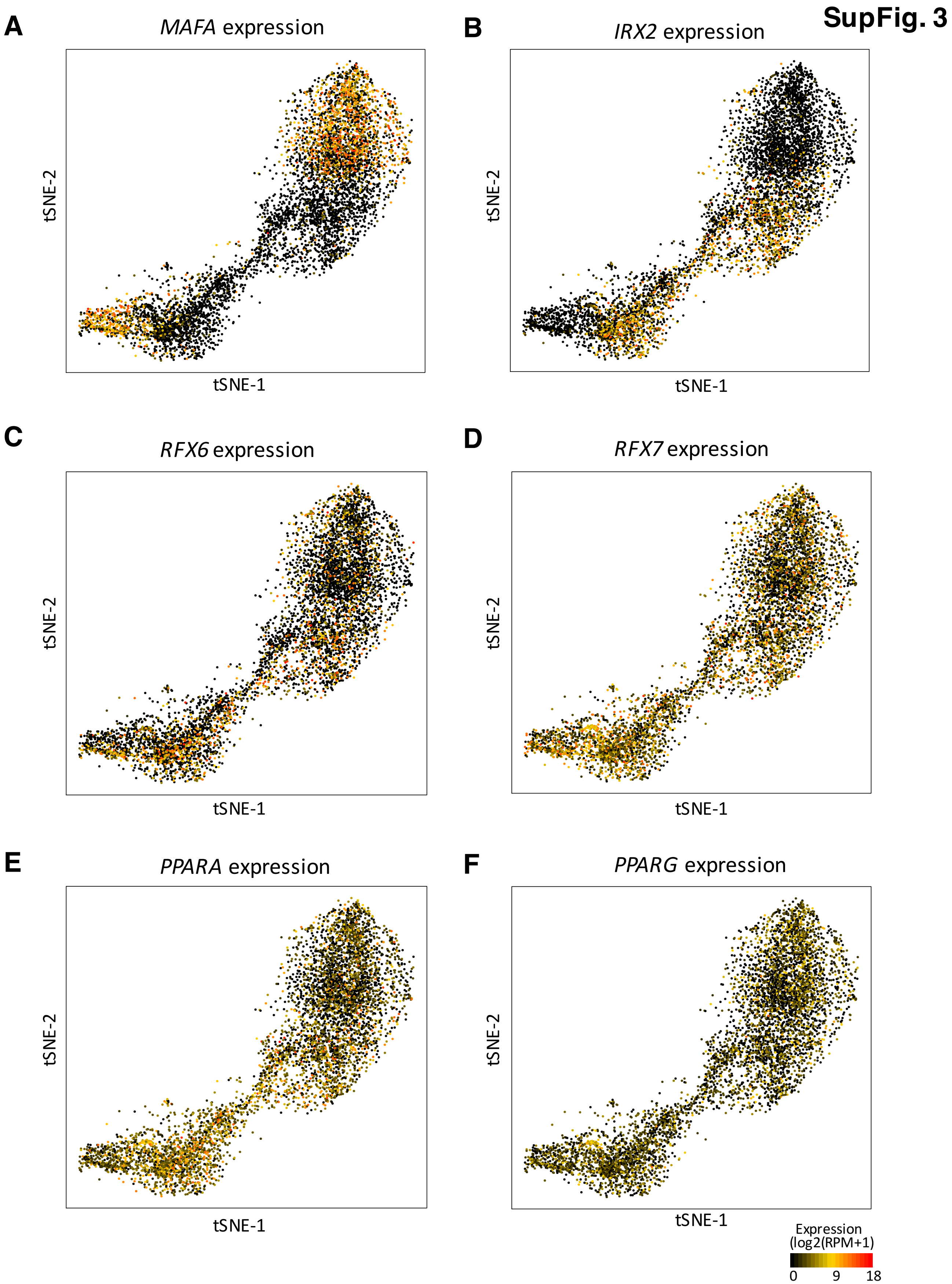


**Supplementary Figure 3. mRNA expression pattern in tSNE plots.** mRNA expression of MafA (A), Irx2 (B), Rfx6 (C), Rfx7 (D), PPARα (E) or PPARγ (F) was visualized as t-SNE plots.

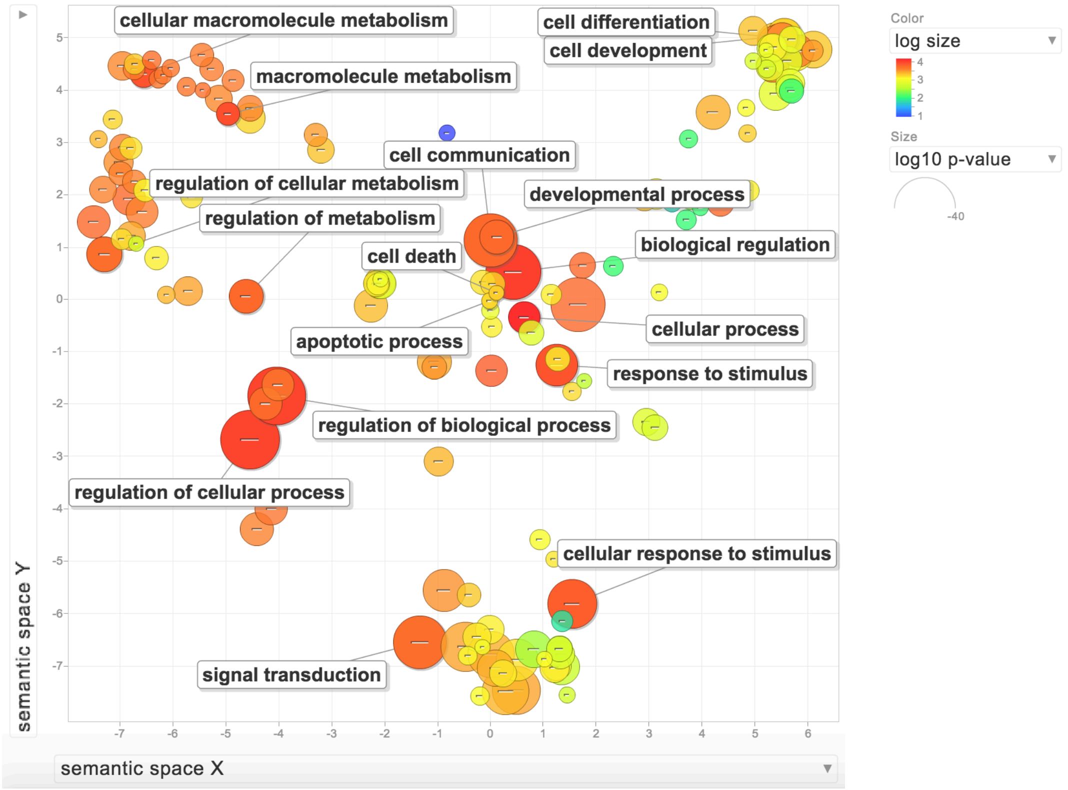


**Supplementary Figure 4. Functional enrichment of GO terms**. Data were analyzed with GO term finder using the list of proteins significantly activated in cluster A compared to cluster B in Figure 1. Functional enrichment was summarized using REVIGO. Enriched terms remaining after the redundancy reduction are represented as scatterplots in a two-dimensional space, which summarizes GO terms' semantic similarities.

**SupFig. 5**

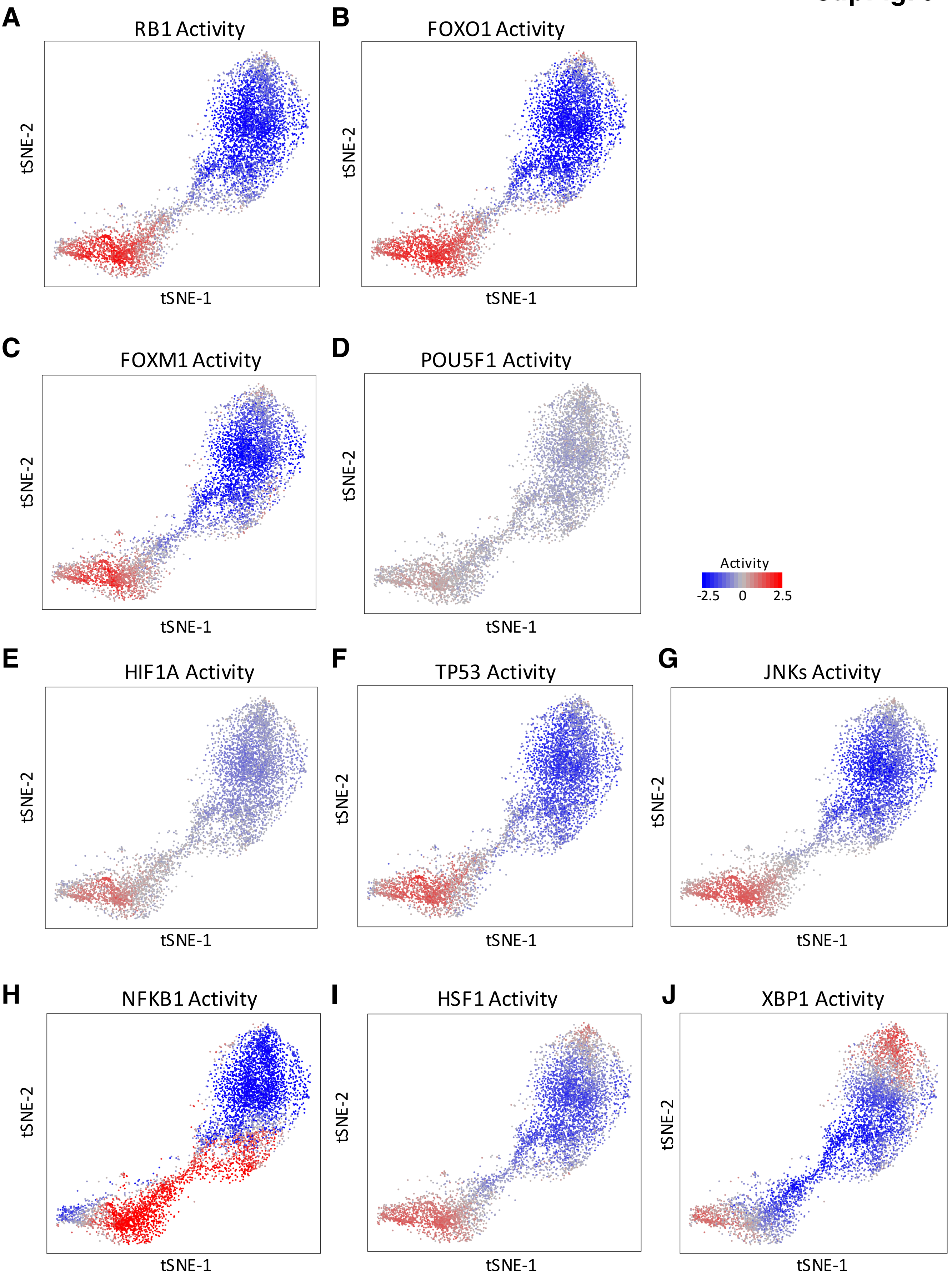

A
RB1 Activity
B
FOXO1 Activity
C
FOXM1 Activity
D
POU5F1 Activity
E
HIF1A Activity
F
TP53 Activity
G
JNKs Activity
H
NFKB1 Activity
I
HSF1 Activity
J
XBP1 Activity
tSNE-1
tSNE-2
Activity
-2.5
0
2.5

**Supplementary Figure 5. Characterization of cells with metabolic inflexibility/stress response and stem-like cell features.** MetaVIPER-inferred activity of metabolic inflexibility regulators (A) RB1 and (B) FOXO1, stemness markers (C) FOXM1 and (D) POU5F1 plotted as t-SNE. Metabolic stress-related transcriptional regulators, including (E) hypoxic stress-related HIF1A, (F) oxidative stress-related TP53, (G) JNK family, (H) NFKB1, (I) HSF1, and (J) ER stress-related XBP1 plotted as t-SNE.

**SupFig. 6**

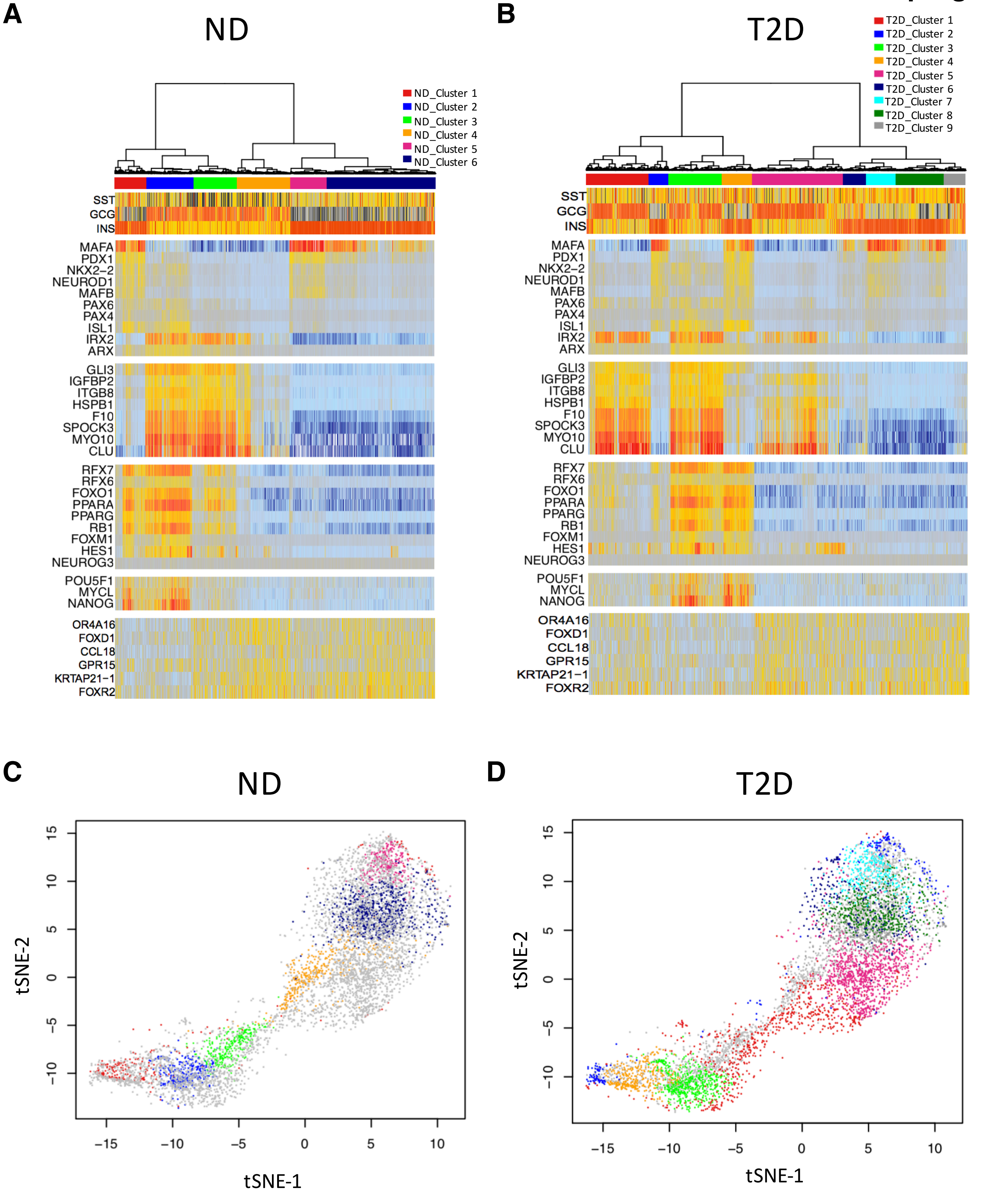


**Supplementary Figure 6. iTerClust classifies human ND or T2D islet cells in different biological states.** (A) hclustering using the iterCluster algorithm structures cell positioning and sub-groups of islet cells only from ND donors. Each sub-group was color-coded accordingly. *SST*, *GCG* and *INS* mRNA expression is plotted at a single-cell level. (B) Same as (A) but from T2D donors. (C) Sub-clusters were projected onto 2D t-SNE space according to metaVIPER-inference for ND islets. Gray dots were a overlay of T2D cells (D) Same as (C) but for T2D islets.

SupFig. 7

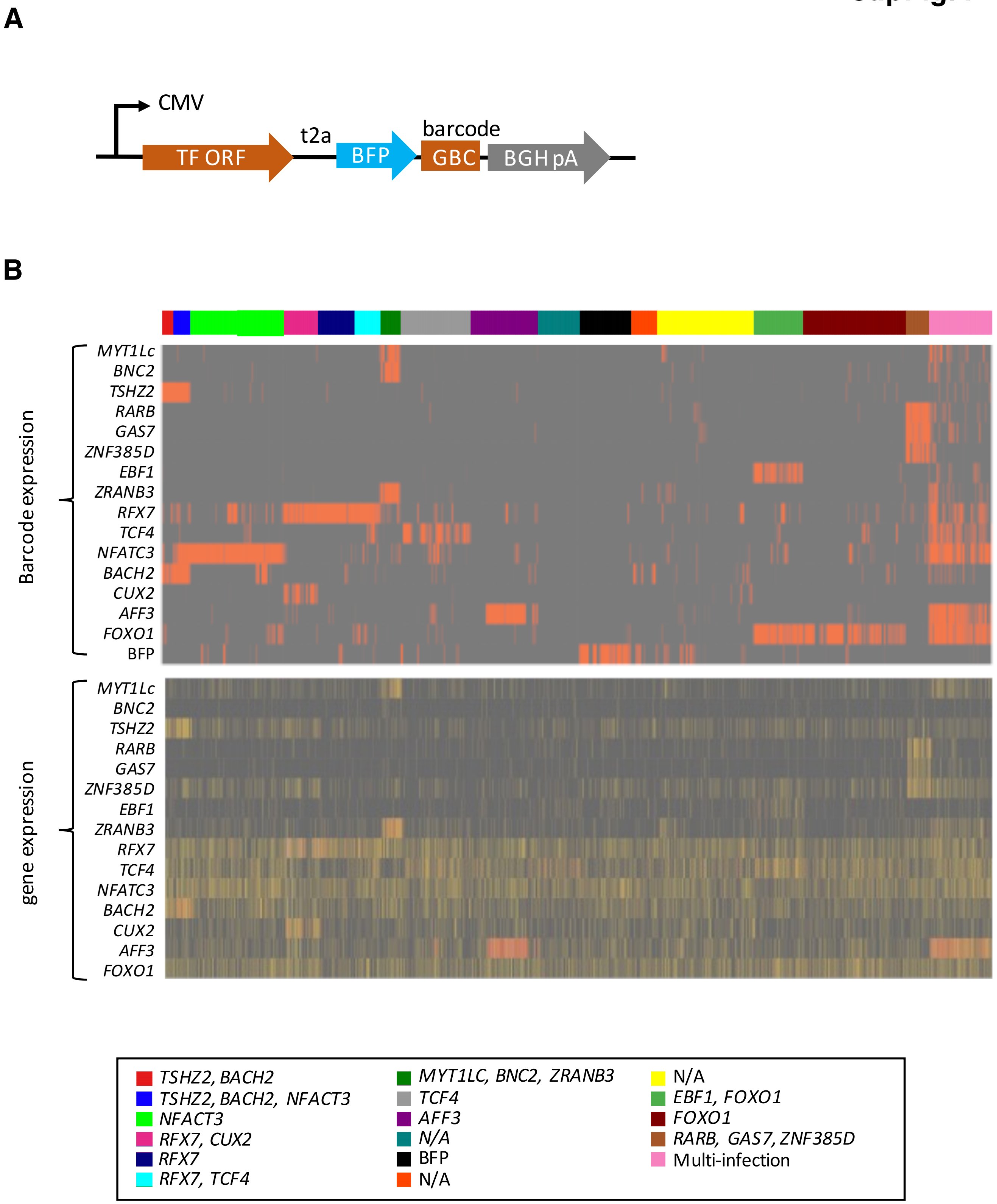


**Supplementary Figure 7. Heatmap to denote cell identity after scGOF-seq.** (A) Schematic drawing of scGOF-seq plasmids. (B) Cell annotation of scGOF-seq is presented for each group, color-coded for single candidate or combinatorial candidate transduction.

**SupFig. 8**

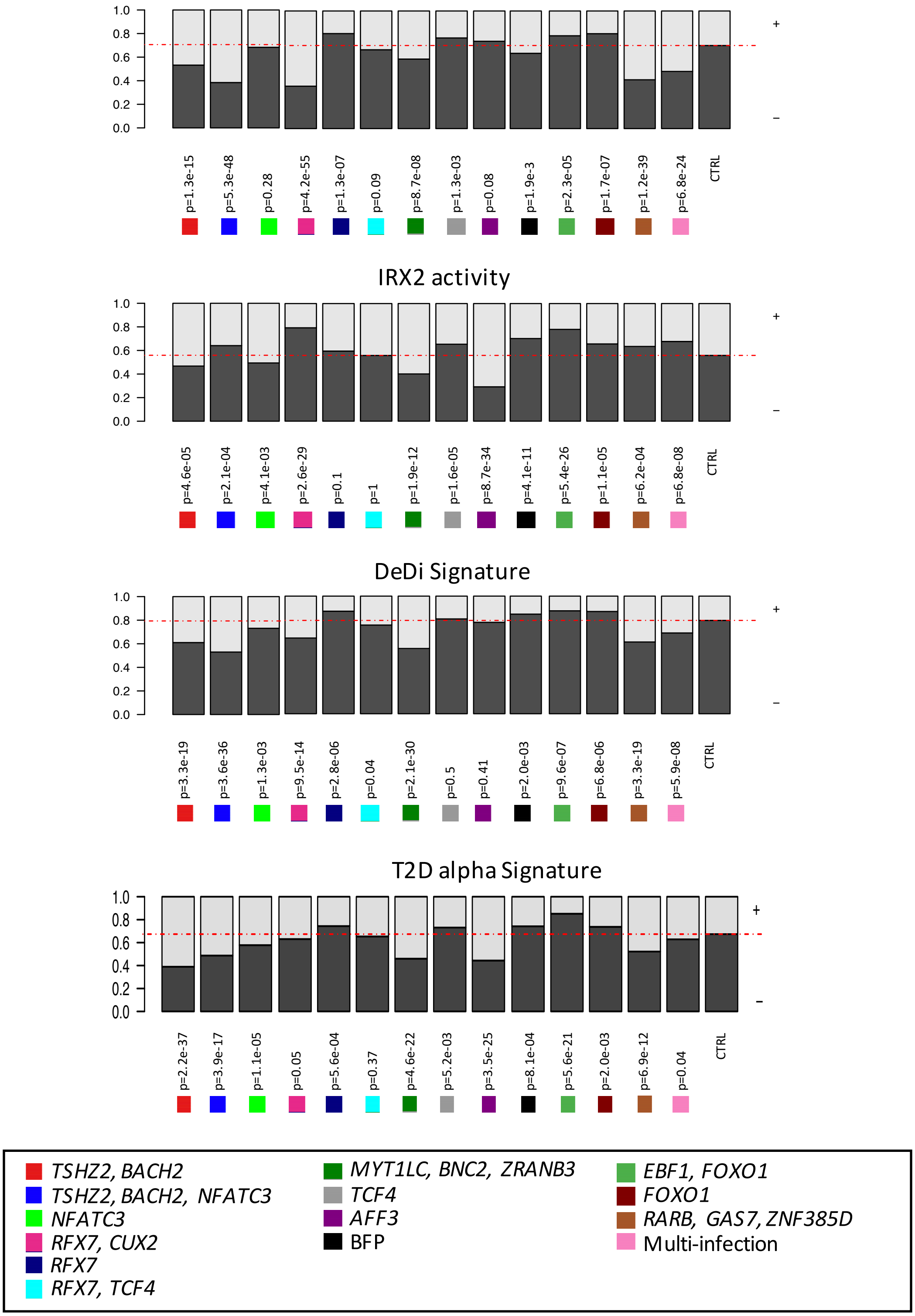


**Supplementary Figure 8. scGOF-seq analyses using ND islets.** (A) Bar-plots showing the proportion of islet cells with positive MAFA activity in each condition. A red dashed line indicates the percentage of islets cells with positive MAFA activity in non-transduced negative controls. (B) Same as (A) but for IRX2 activity. (C) Bar-plots showing the proportion of islet cells with a positive DeDi signature (> activity 0) in each condition. (D) Bar-plots showing the proportion of islet cells with a positive B6-like signature (> activity 0) in each condition.

**SupFig. 9**

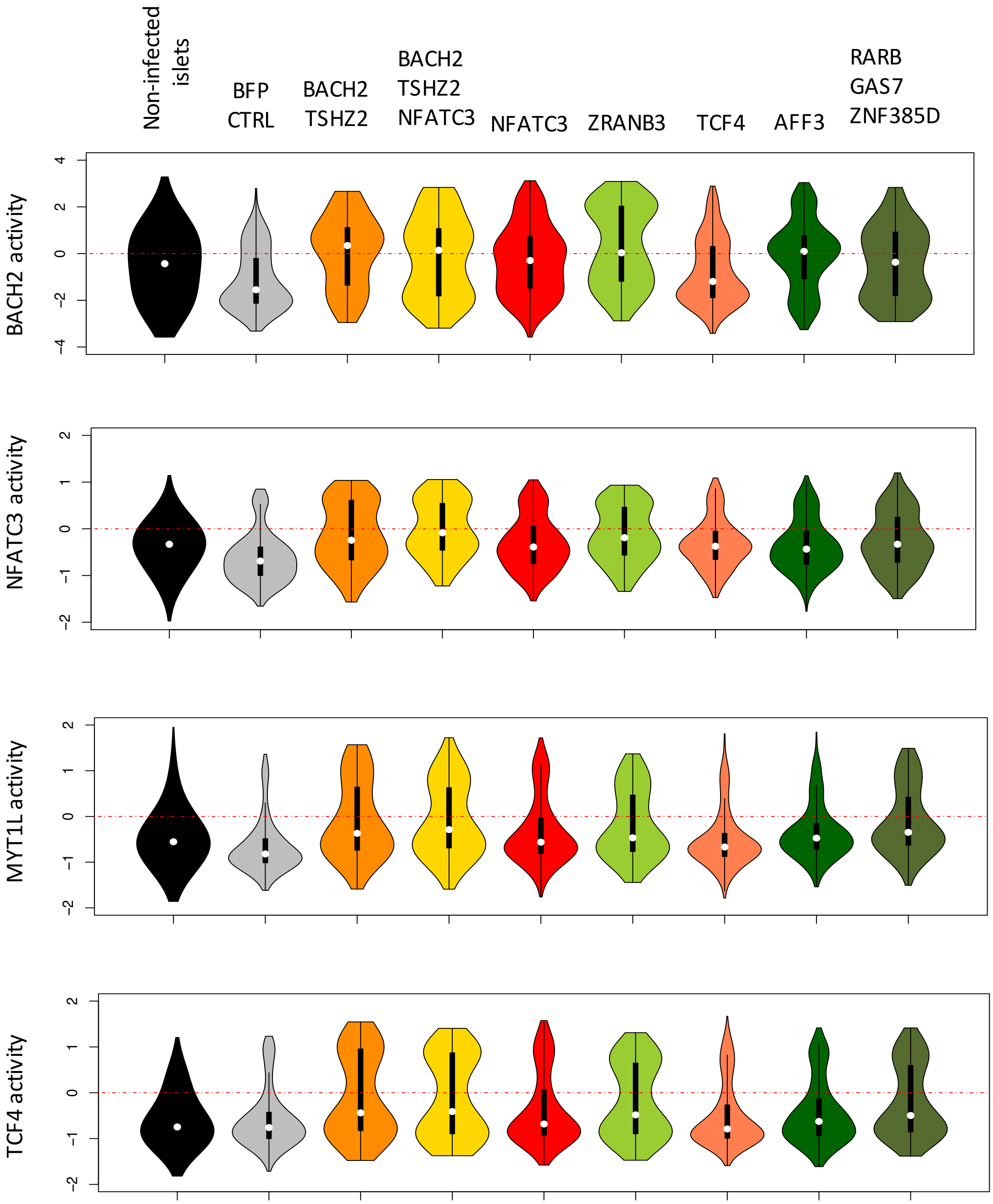


**Supplementary Figure 9. Core driver network of ND cell conversion into DeDi signature cells.** Violin plots of core driver network, BACH2, NFATC3, MYT1L and TCF4 for each single candidate or in combination according to DeDiSig score.

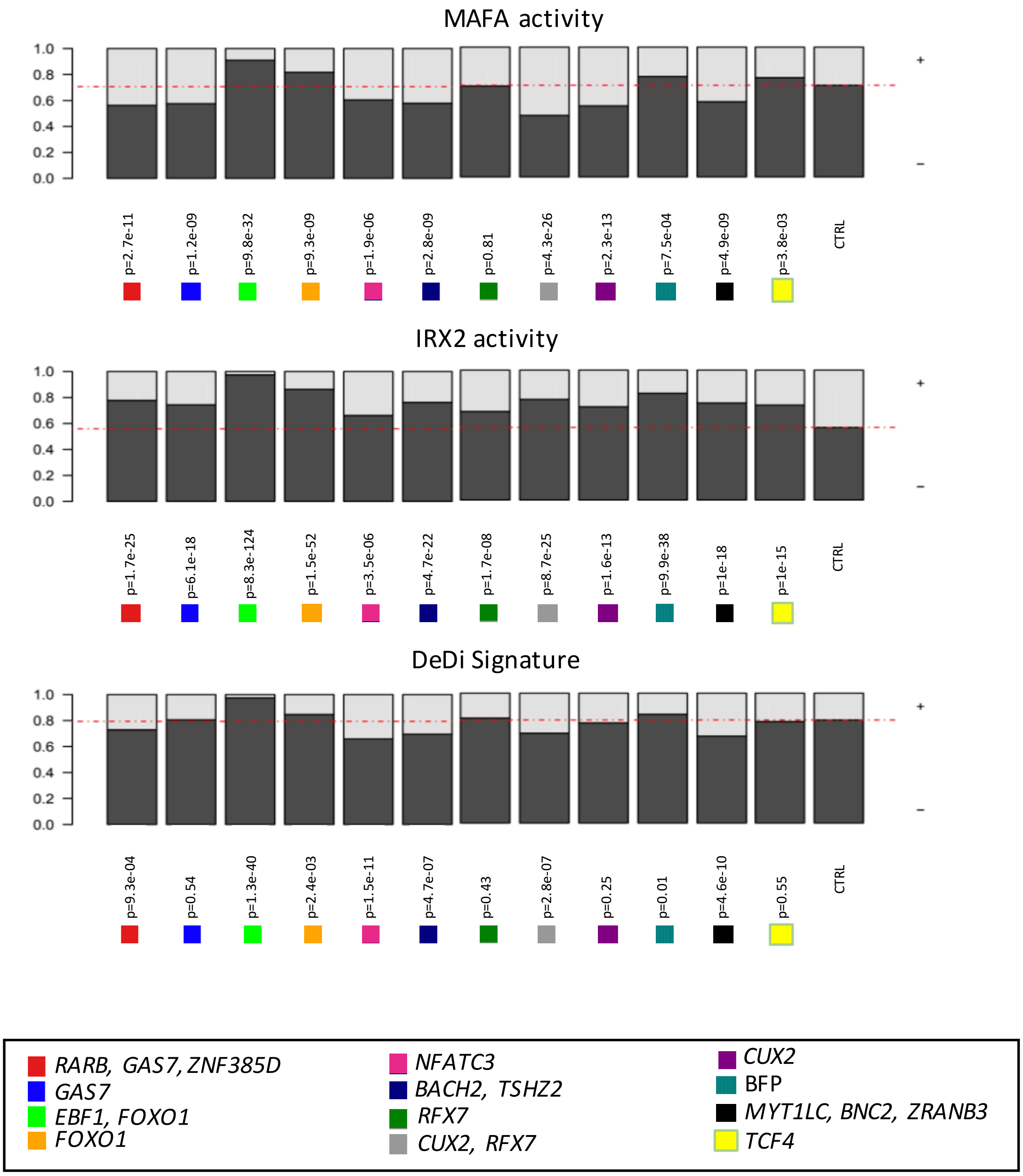


**Supplementary Figure 10. Biological scGOF-seq replicate.** (A) Bar-plots showing the proportion of islet cells with positive MAFA activity in each gain-of-function condition. A red dashed line indicates the percentage of islets cells with positive MAFA activity in non-transduced negative control. (B) Same as (A) but for IRX2 activity. (C) Bar-plots showing the proportion of islet cells with a positive DeDi signature (> activity 0) in each condition.
